# Developmental Stage-Dependent Transcriptomic Responses to Neonatal Intraventricular Hemorrhage

**DOI:** 10.1101/2025.07.11.664369

**Authors:** Elizabeth Wallace-Anthony, Miriam Zamorano, Hemendra J Vekaria, Braden B Oldham, Kiara P Umpornpun, Scott D Olson, Stefano Berto, Brandon A Miller

## Abstract

Neonatal intraventricular hemorrhage (IVH) is a major complication of preterm birth, yet how developmental stage influences the brain’s response to injury remains unclear. We performed single-nucleus RNA sequencing on rat brains 24 hours after IVH at postnatal day 2 (PND2) or day 5 (PND5) to define transcriptional responses across cell types. We identified 42 distinct cell populations and found that PND5 brains exhibited a markedly stronger immune and inflammatory response to IVH, with a threefold increase in differentially expressed genes compared to PND2. Microglia were the most perturbed cell type at both stages, showing increased oxidative stress and polarization toward both pro- and anti-inflammatory phenotypes at PND5. Ligand-receptor and regulon analysis revealed a shift from reparative IGF2 and TGF-β signaling at PND2 to proinflammatory Wnt signaling and activation of Runx1 and Stat5 at PND5. These findings highlight the importance of developmental timing in shaping the neuroimmune response to IVH and identify potential stage-specific therapeutic targets.

## INTRODUCTION

Neonatal intraventricular hemorrhage (IVH) is a common complication of preterm birth, especially in preterm infants with low or very low birth weight. IVH originates from the underdeveloped germinal matrix, a site of rapid cell division adjacent to the lateral ventricles of the brain. The germinal matrix is highly vascular and is vulnerable to hemodynamic instability and inflammation related to premature birth [1]. When fragile blood vessels within the germinal matrix rupture, blood is released into the brain parenchyma and the lateral ventricles. Children who experience IVH have persistently high levels of blood breakdown products in their intraventricular cerebrospinal fluid (CSF) for weeks after the ictus. This is visible on cranial ultrasounds performed after IVH and is documented in clinical studies of CSF from children with IVH [2, 3]. The persistence of blood products in the CSF after IVH allows these factors to interact with the subventricular zone and penetrate throughout the brain parenchyma via an injured CSF-ependymal barrier [4]. During the same period in which IVH pathology develops, the brain undergoes rapid development, including synaptogenesis, myelination, and development of the neuroimmune system [5].

Many animal models are available for studying IVH. Some studies use collagenase, which induces a primarily destructive lesion with a secondary release of blood into the ventricles [6]. Others use intraventricular injections of blood or blood products to model the effects of a gradually resolving blood clot within the ventricular system [7]. These two experimental models have been compared in adult animals and have shown significant overlap in the transcriptional profiles they produce [8]. Our recent work testing intracerebroventricular blood delivered at postnatal day (PND) 2 and PND5 in separate cohorts of animals revealed that the immune reaction to IVH differed greatly by developmental stage and that white matter was only affected in animals that underwent IVH at PND5 but not at PND2 [9]. This prior study relied on flow cytometry to differentiate the innate immune response at different developmental stages. However, the entire landscape of the central nervous system (CNS) is rapidly changing during this developmental window [5]. IVH outcomes are likely determined by a combination of factors including the changing neuroimmune system and the other cells’ changing vulnerability to injury based on developmental stage [10, 11].

This study was designed to compare the differential responses of the developing CNS to intraventricular blood components and to reveal the behavior of all CNS cell types via single-nuclei RNA analysis. It was also designed to analyze sex as a biological variable in response to IVH. These results help paint a more complete picture of how CNS development affects the response to IVH. Our analysis focused on neuroimmune reactions to injury, in an attempt to highlight how CNS development affects the inflammatory response to IVH and vice versa. Our data indicate that the PND2 and PND5 brains differ in cellular composition and reaction to intraventricular blood products, with the PND5 brain exhibiting a more robust expression of immune relevant genes after IVH. This dataset will allow the generation of many testable hypotheses and identification of therapeutic targets in neonatal IVH.

## METHODS

### Overall Distribution and design

Rats underwent IVH at postnatal day 2 (PND2) and postnatal day 5 (PND5), and their brains were harvested for analysis 24 hours later (Figure 1A) [9]. The experiment used 32 rats overall, with 16 for each timepoint. Of these 16, 8 were control and 8 underwent IVH at each timepoint. Four of each set were male and the other 4 were female (Supplementary Figure 1A).

**Figure 1.**
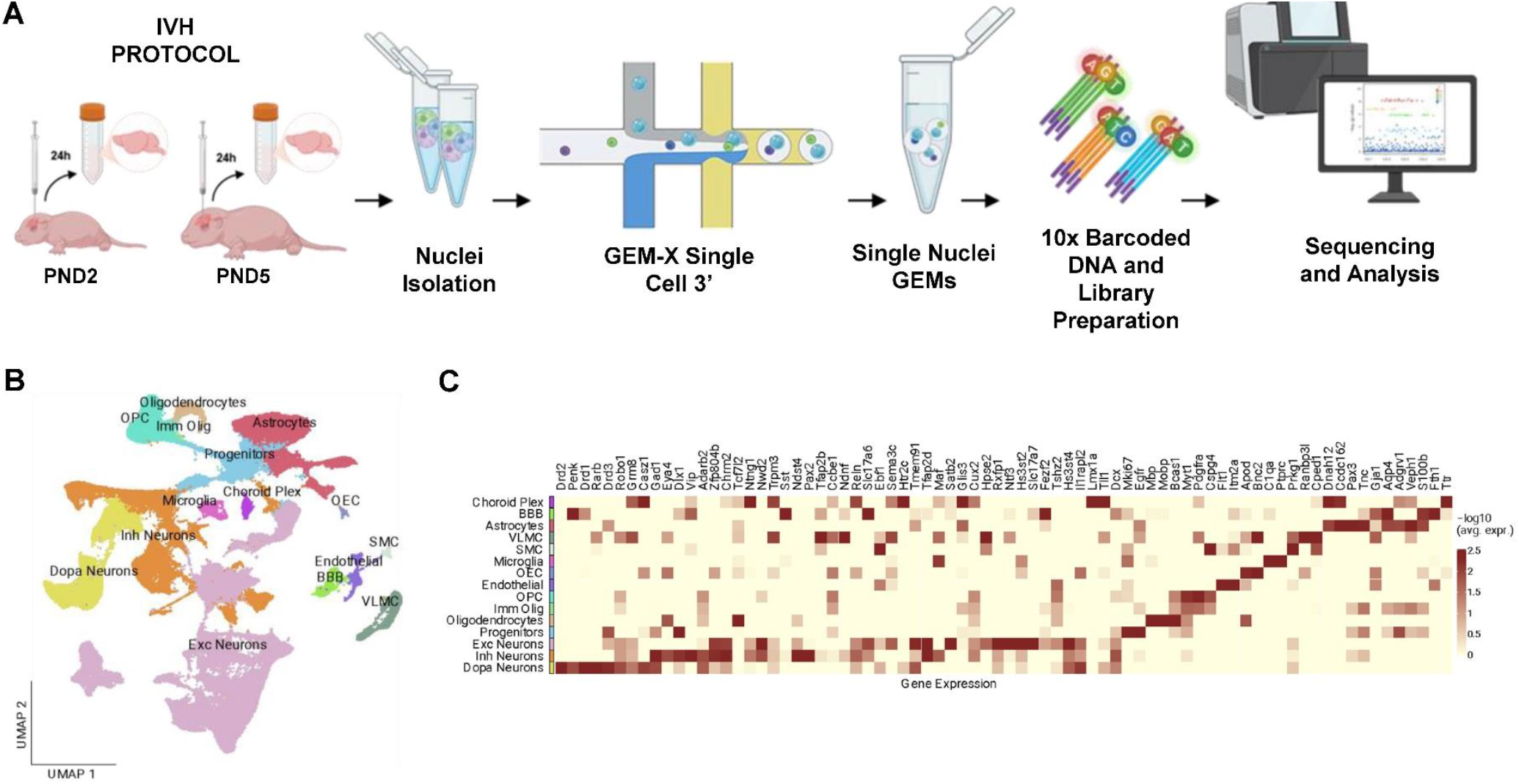
Experimental Design and Analysis Workflow. Experimental design. Rats were subjected to IVH and the brains harvested 24 hours later. Nuclei were then isolated and loaded onto Chromium GEM-X 3’ Chip. Libraries were then sequenced and analyzed. **B**. UMAP depicting the aggregated data from both IVH and Control rats with cell annotation. **C**. Heatmap showing scaled gene expression of marker genes for each cell type.

### Intraventricular injections

All animal protocols were approved by our Institutional Animal Care and Use Committee. Intraventricular injections were performed as we have previously described [9]. Timed pregnant Sprague Dawley rats from Charles River Laboratories (Wilmington, MA, USA) were housed in a controlled temperature room under a 14/10-hour light/dark cycle with food and water ad libitum. Rat pups were separated by sex and postanal day PND2 and PND5 groups for intraventricular injection. For injections, the scalp was prepped with betadine and a 30G needle was placed directly through the scalp with the following coordinates: for PND2, 3.8 mm from lambda, lateral 0.7 mm, 2 mm deep, and for PND5, 4.6 mm from lambda, lateral 1.1 mm, and 3.3 mm deep, measured through transillumination of the cranial sutures. 30μl of lysed blood from a female donor was injected into the right ventricle at a flow rate of 0.5 μl/sec. 30μl of sterile artificial CSF (Biotechne Tocris Cat. 3525) was used as a control. The needle was left in place for 1 min after the injection to prevent retrograde flow out of the syringe tract. After the injections, the animals were immediately returned to their home cage and dam for recovery.

### Brain Extraction

24h after IVH, animals were euthanized by rapid decapitation and meningeal tissue was removed. Whole brains were placed into 2 ml Eppendorf tubes in liquid nitrogen for fast freezing.

### Nuclei Isolation

Brains were placed on ice with lysis buffer Tris-HCl (pH 7.4) 10mM, NaCl 10 mM, MgCl23mM, Nonidet P40 Substitute (MilliporeSigma Cat. 74385) 0.1%, in nuclease-free Water) for 15 minutes. Using a 2ml pipette, the tissue was gently transferred to a 50ml falcon tube adding 5ml of Hibernate E/B27/GlutaMAX (HEB) medium (Gibco Cat. A12476-01). The tissue was triturated with a pasteur pipette and the generated homogenates were strained with 30 μm MACS SmartStrainer to remove large debris and centrifuged at 500 RFC for 5 min at 4°C, supernatant was removed and 10 ml of nuclei washing solution and resuspension buffer 1% BSA (MilliporeSigma Cat. 3335399001) in 1X PBS was added to the pellet. Lysis efficiency and viability were assessed by staining cells with trypan blue and counted using the Countess II FL Automated Cell Counter/ microscopy.

### Single-cell RNA sequencing

Isolated nuclei were processed by UTHealth Cancer Genomics Core at The University of Texas Health Science Center at Houston (CPRIT RPND240610). Single nuclei capture and library construction were performed by following the Chromium GEM-X Single Cell 3’ Reagent Kits v4 protocol (CG000731). Briefly, cells were resuspended in PBS with 0.04% BSA and RNase inhibitor at a concentration of 700-1200 cells/µl. The single nuclei were loaded onto Chromium GEM-X 3’ Chip (PN-1000690) with partitioning oil and barcoded single-cell gel beads. The barcoded and full-length cDNA was produced after incubation of gel beads-in-emulsion (GEMs) and amplified via PCR for library construction. The library preparation was performed by following the protocol of Chromium GEM-X Single Cell 3’ Kit v4 (PN-1000691). The quality of the final libraries was examined using Agilent High Sensitive DNA Kit (#5067-4626) by Agilent Bioanalyzer 2100 (Agilent Technologies, Santa Clara, USA). The qualified libraries were sent for the paired-end sequencing on an Illumina Novaseq System (Illumina, Inc., USA).

### Single nucleus RNA sequencing Analysis

Reads were aligned to the rat mRatBN7 reference genome with introns from UCSC using the 10x Genomics CellRanger analytical pipeline [12, 13]. CellRanger was used to perform dataset aggregation to normalize for total numbers of confidently mapped reads using the “count” function to generate raw gene-cell expression matrices. CellBender was used to infer and remove ambient RNA contamination [14]. Data was then combined and integrated with the *Seurat* package (v5.2.1) in R into a Seurat object using the Read10X function to import the data [15]. Replicate information was integrated into the Seurat metadata and replicates were verified to have similar distributions of cell type in each replicate (Supplemental Figure 1A). Filtering was also performed to remove cells with mitochondrial content over 5 percent, less than 25000 unique molecular identifiers (UMIs), nuclear fraction (NF) greater than 0.5, and total gene numbers under 250 (Supplementary Figure 1B-D). Doublets were also identified and removed using *scDblFinder* (v1.18.0) [16]. Data was normalized to account for variation between replicates using *SCTransform*, which was utilized for integration and to identify highly variable genes [17]. Principal components analysis (PCA) was performed on a selection of the 3000 most variable genes, and the top 30 significant principal components (PCs) with a resolution of 0.3 for Louvain clustering were selected for Uniform Manifold Approximation and Projection (UMAP) and visualization of gene expression. Data was then integrated with *Harmony* (v 1.2.3) using the 30 PCs and 3000 most variable genes [18]. Cluster marker genes were identified using FindAllMarkers with the Wilcoxon Rank Sum test using standard parameters (Figure 1B-C, Supplemental Figure 1E-I). Cell annotation was then performed manually using gene markers by clustering (Figure 1B-C, Supplemental Figure 1E). Cell annotation wes further confirmed with a single nuclei reference data of whole mouse brain [19].

### Perturbation analysis

Cell proportion inference by experimental condition was performed using R package *miloR* (v2.3.1) [20]. The buildGraph, makeNhoods, testNhoods, and calculateNhoodDistance functions were used in sequence to group cells into neighborhoods and calculate how the cell counts changed in the neighborhoods between experimental conditions, with data split by timepoint. The plotNhoodGraphDA function was then used in combination with the *Seurat* (v5.2.1) FeaturePlot function to plot results [15]. Cell type prioritization by experimental perturbation was performed using Python package meld (v1.0.2) [21]. The fit_transform function was used to prepare the data and the normalize_densities was used to calculate perturbation likelihood by experimental condition with data split by time point. Data was plotted in R using *ggplot2* (v3.5.1). Verification of MELD results was performed using R package Augur (v1.0.3) [22]. The calculate_auc function was used to determine AUCs of experimental perturbation by experimental condition, with data split by time point. These scores were then ranked and plotted using *ggplot2* (v3.5.1) and *ComplexHeatmap* (v2.20.0) [23].

### Identification of differentially expressed genes

To identify differentially expressed genes (DEGs) between IVH-induced rats and control at each time point, we performed a pseudobulk analysis within each cell subtype. Briefly, raw counts were aggregated by sample using the R function AggregateExpression in Seurat. The resulting pseudobulk count matrices, one for each cell type, were analyzed using the DESeq2 package (v1.44.0), which fits a Gamma-Poisson generalized linear model [24]. Number of nuclei per cell type was included as technical covariates in the model. Genes were defined as significantly differentially expressed at FDR-adjusted p-value<0.05. Differentially expressed genes (DEGs) were visualized and plotted using *ggplot2* (v3.5.1) and *ComplexHeatmap* (v2.20.0) [23]. Overlapping DEGs by cell type were also calculated and plotted as an upset plot using *ggplot2* (v3.5.1). A list of immune-relevant genes was used to highlight immune-relevant DEGs. Gene fold changes and significant DEGs were plotted on KEGG pathway maps for pathways of interest using *ggkegg* (v1.4.1) [25-27].

### Functional enrichment

Functional enrichment for immune pathways was performed on each subset of cell types using *AUCell* (v1.26.0) [28]. For each timepoint, RNA count data from the Seurat object was used as the expression matrix input. Immune pathways were acquired for a list of immune relevant genes using getGmt from *GSEAbase* (v1.66.0) [29]. Area under the curve (AUC) values were calculated for each pathway by cell type using AUCell_run and AUCell_calcAUC functions. A wilcoxon t-test using *rstatix* (v0.7.2) with falsediscovery rate (FDR) adjustment was then performed between control and IVH using AUC values for each pathway and cell type. Differences in AUC between control and IVH were then ranked and plotted with significance annotated using *ggplot2* (v3.5.1).

### Cell-cell communication analysis

Intercellular communication analysis was performed across all cell types with microglia as the target cell type of interest using *nichenetr* (v2.2.0) [30]. The mouse ligand receptor network and ligand target matrix were used due to the similarity in genomes. The get_expressed_genes function was used to extract expressed genes from the microglia as the receiver and all cell types as senders and these were used as inputs for the predict_ligand_activities function to predict ligand interactions. The top ranked ligands and receptors were then filtered for and graphed using the make_heatmap_ggplot function.

### Cell-type specific eRegulons

Transcriptomic regulons were inferred using the pySCENIC pipeline (v0.12.1+8.gd2309fe) [28, 31]. Seurat objects were first stratified by developmental time point and subsequently by treatment conditions. Each subset was converted to AnnData and loom formats, followed by sequential execution of the GRN, CTX, and AUCELL steps in pySCENIC to generate condition-specific regulon activity matrices in loom format. The resulting output files were imported into Python for post-processing, and the regulon activity scores were exported as CSV files. These data were then integrated with the corresponding Seurat objects in R for downstream visualization. The top ten regulons in microglia were identified and visualized for both control and IVH conditions using ggplot2 (v3.5.1) and ComplexHeatmap (v2.20.0) [23].

### Macrophage polarization analysis

Macrophage polarization analysis of microglia was conducted using MacSpectrum (v1.0.1), employing the macspec function to calculate polarization indices [32]. Statistical comparisons of the Macrophage Polarization Index (MPI) across time points and treatment conditions were performed using the Wilcoxon rank-sum test with false discovery rate (FDR) correction, implemented via the rstatix package (v0.7.2). Resulting significance levels and group differences were visualized using ggplot2 (v3.5.1). Additionally, in Python, seaborn (v0.13.2) was used to generate joint plot visualizations of MPI and the Activation-induced Macrophage Differentiation Index (AMDI) for microglia in the IVH condition across all time points. It is important to note that MacSpectrum was originally developed and optimized for macrophages in human and mouse datasets, which may limit its sensitivity to microglial-specific or species-specific polarization states in our model system.

### Code Availability

Figures and supplementary tables were generated using custom scripts in R (v4.2.1) and Python. The code used for data analysis in this study is available on GitHub: https://github.com/BioinformaticsMUSC/Rat_IVH_SingleCellAnalysis

## Data availability

All raw IVH snRNA-seq data are deposited to the GEO archive under the accession number: GSE302024.

The data presented in this manuscript are available through an interactive Shiny application. These applications include data from IVH PND2 and IVH PND5 datasets: https://bioinformatics-musc.shinyapps.io/Miller_Whole_Brain_PND2;https://bioinformatics-musc.shinyapps.io/Miller_Whole_Brain_PND5

## RESULTS

### Single-Nucleus RNA Sequencing Defines Major Cell Types and Subclasses in the Neonatal Rat Brain Following IVH

To understand how IVH affects the cellular composition of the neonatal brain, we first defined the major cell types present using single-nucleus RNA sequencing. Using the single-nucleus RNA sequencing (snRNA-seq) platform from 10x Genomics, we obtained transcriptomic data from 161,933 nuclei from 32 rat brains (Figure 1A, Supplementary Figure 1A-B). After filtering out doublets and low-quality nuclei, 129,279 high-quality nuclei remained for downstream analysis (Supplementary Figure 1B-D). Each nucleus yielded a mean of 1409 genes and 2890 unique molecular identifiers (UMIs) (Supplementary Figure 1B-D).

Clustering analysis revealed 15 major cell populations of the rat brain (Figure 1B). Cell type annotation was guided by known marker gene expression patterns from non-injured mouse cortices (Figure 1C, Table S1) [19]. Our analysis identified four major cell classes: glutamatergic/excitatory (Exc; 52,367 nuclei), GABAergic/inhibitory (Inh; 20,114 nuclei), dopaminergic (Dopa; 12,349 nuclei), and glial (Glia; 43,449 nuclei), which were further subdivided into 42 transcriptionally distinct subclasses (Supplementary Figure 1E). These included 12 excitatory neuronal clusters (Supplementary Figure 1F),

10 inhibitory neuronal clusters (Supplementary Figure 1G), 4 dopaminergic neuronal clusters (Supplementary Figure 1H), and 16 glial clusters (Supplementary Figure 1I).

### Developmental Stage Influences Cell Type Composition and Perturbation more so than IVH

Next, we analyzed differences in cell composition between IVH and controls, focusing on the 42 cell subtypes annotated at high resolution and different time points. We performed differential abundance analysis to compare cell proportions in the control and IVH groups within local neighborhoods using the Milo k-nearest neighbor (k-NN) approach [20]. The overall distribution of cell types remained relatively consistent between control and IVH groups (Figure 2A). In contrast, the developmental stage had a pronounced impact on cell type distribution, with changes at PND5 reflecting maturation of various cell types (Figure 2B). Specifically, there was a marked increase in oligodendrocytes, oligodendrocyte precursor cells (OPCs), olfactory ensheathing cells (OECs), and microglia, accompanied by a reduction in blood-brain barrier (BBB) cells and choroid plexus cells relative to PND2 (Figure 2B). Additionally, specific astrocyte and neuronal subtypes showed increased representation at PND5 in the IVH condition (Supplementary Figure 2A), likely due to increased neuronal activity and gene expression in response to IVH at this time point. Notably, across both developmental stages, most cell types exhibited minimal fold changes between spatial neighborhoods when comparing control and IVH groups (Figure 2A–B). An exception was observed at PND5, where a specific excitatory neuronal subtype (cerebral anterior hypothalamus glutamatergic neurons (CNU HYA Glut)) showed a distinct increase in representation, again likely due to increased activity and gene expression (Figure 2B, Supplemental Figure 1D).

**Figure 2.**
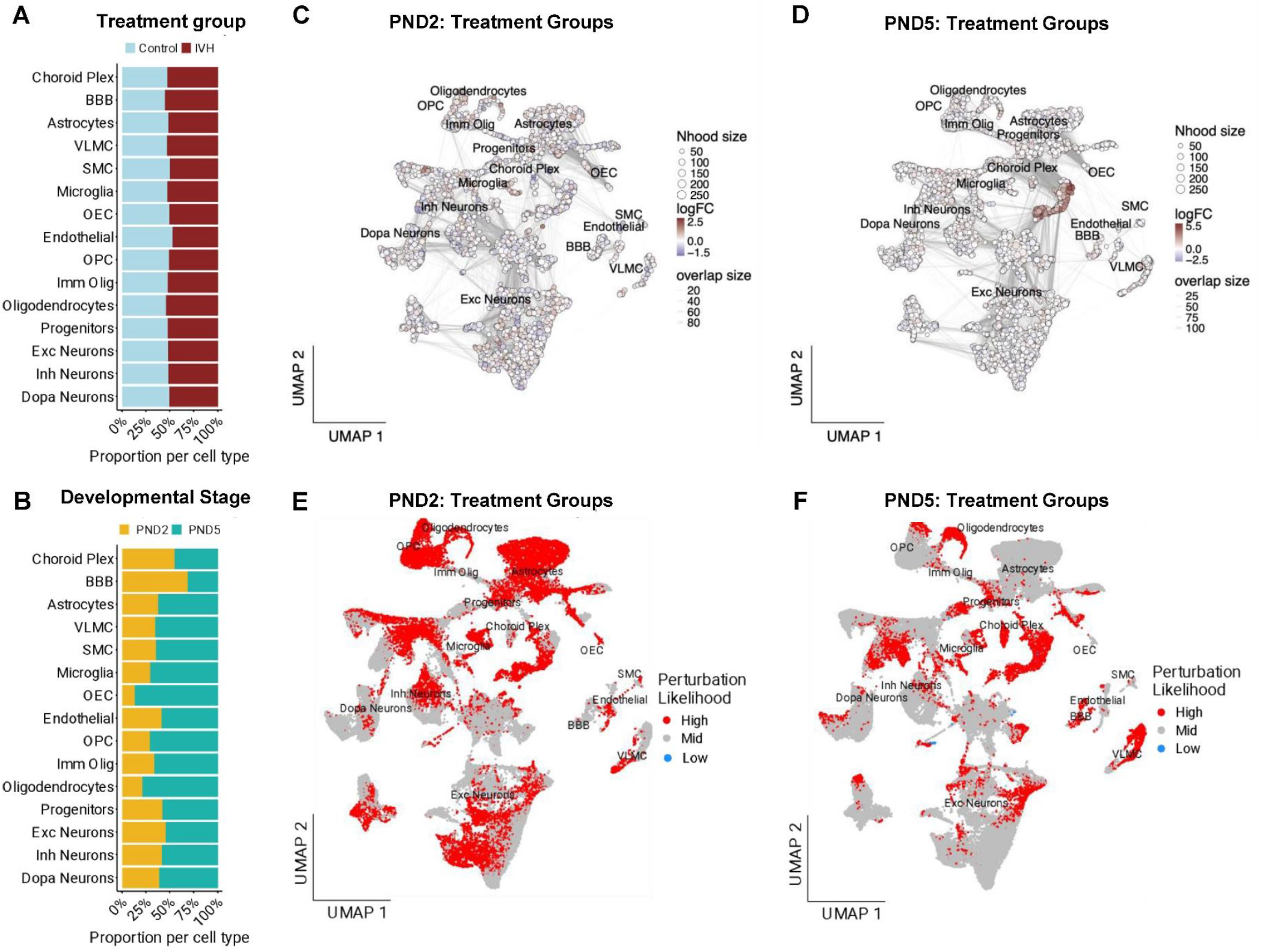
Cell perturbation following IVH. **A**. Stacked barplot showing the proportion explained by IVH and control samples for each cell type. **B**. Stacked barplot showing the proportion explained by developmental stage. **C-D**. Differential abundance analysis comparing the abundance in IVH and control rats for PND2 (C) and PND5 (D). **E-F**. UMAP of annotated cell types showing MELD likelihoods for IVH compared with control in PND2 (E) and PND5 (F).

Comparing perturbation for each timepoint affected by control and IVH using MELD, a graph-based method that quantifies changes in local cellular neighborhoods, we observed that the overall high-perturbation cell types were largely similar, although astrocytes, and OPCs had more perturbation at PND2, as did microglia, although some microglia still had high perturbation at PND5 (Figure 2E-F) [21]. Most cell types had significant perturbation after IVH at both time points (Supplementary Figure 2B-C). These results were corroborated by Augur, a machine-learning approach that prioritizes cell types based on their transcriptional distinction between conditions, which yielded consistent findings (Supplementary Figure 2D-F).

### Transcriptomic Analysis Reveals Enhanced Inflammatory and Adaptive Gene Expression Responses to IVH at PND5

To further elucidate the molecular impact of IVH on different cell types, we performed differential gene expression analysis comparing IVH and control groups at PND2 and PND5 using a pseudobulk approach based on DESeq2. We identified 99 differentially expressed genes (DEGs) overall at PND2 and 388 DEGs overall at PND5, suggesting that IVH has a greater effect on gene expression at PND5 (Figure 3A-B, Table S2). PND5 had over three times the number of DEGs at PND2 across more cell types. Microglia had the greatest number of DEGs overall, and of these, 13 were immune-relevant genes at PND2 and 40 were immune-relevant at PND5 (Figure 3 A-B, Supplemental Figure 3A-B). When gene ontology enrichment was performed for each category of cell type, the most commonly enriched immune-relevant pathway with significant changes between control and IVH was oxidative phosphorylation at PND5 (Figure 3C-F). No pathways showed significant changes at PND2. In glial cells, the most activated pathway is the reactive oxygen species pathway. This correlates with our prior work showing the sequelae of oxidative stress concentrated in white matter in a hemoglobin-injection model of neonatal IVH [33]. Microglial Abundance and Polarization in Response to IVH. As microglia are the cell type most affected by IVH, we also examined how their abundance changed across conditions. Microglial numbers significantly increased from PND2 to PND5 in both control and IVH groups. However, a significant difference between control and IVH was observed only at PND2, with IVH animals exhibiting more microglia (Figure 3G). By PND5, overall microglial numbers were comparable between groups, yet microglia in the IVH condition showed a markedly elevated number of DEGs and a shift toward both proinflammatory (M1-like) and reparative (M2-like) polarization states (Figure 3H-J, Figure 4). Although a binary “M1 versus M2” microglial state is an oversimplification, and M1 and M2 phenotypes can coexist in pathology, we adopted this framework to describe relative polarization dynamics [34]. Notably, IGF-1 related genes were significantly downregulated after IVH in PND2 animals but not in PND5. Given that IGF-1 signaling by microglia promotes white matter development, its downregulation at PND2 may contribute to IVH-induced hypomyelination [35]. In PND5 microglia, CD44 was highly upregulated. This toll-like receptor has been associated with the progression of inflammatory diseases [36, 37].

**Figure 3.**
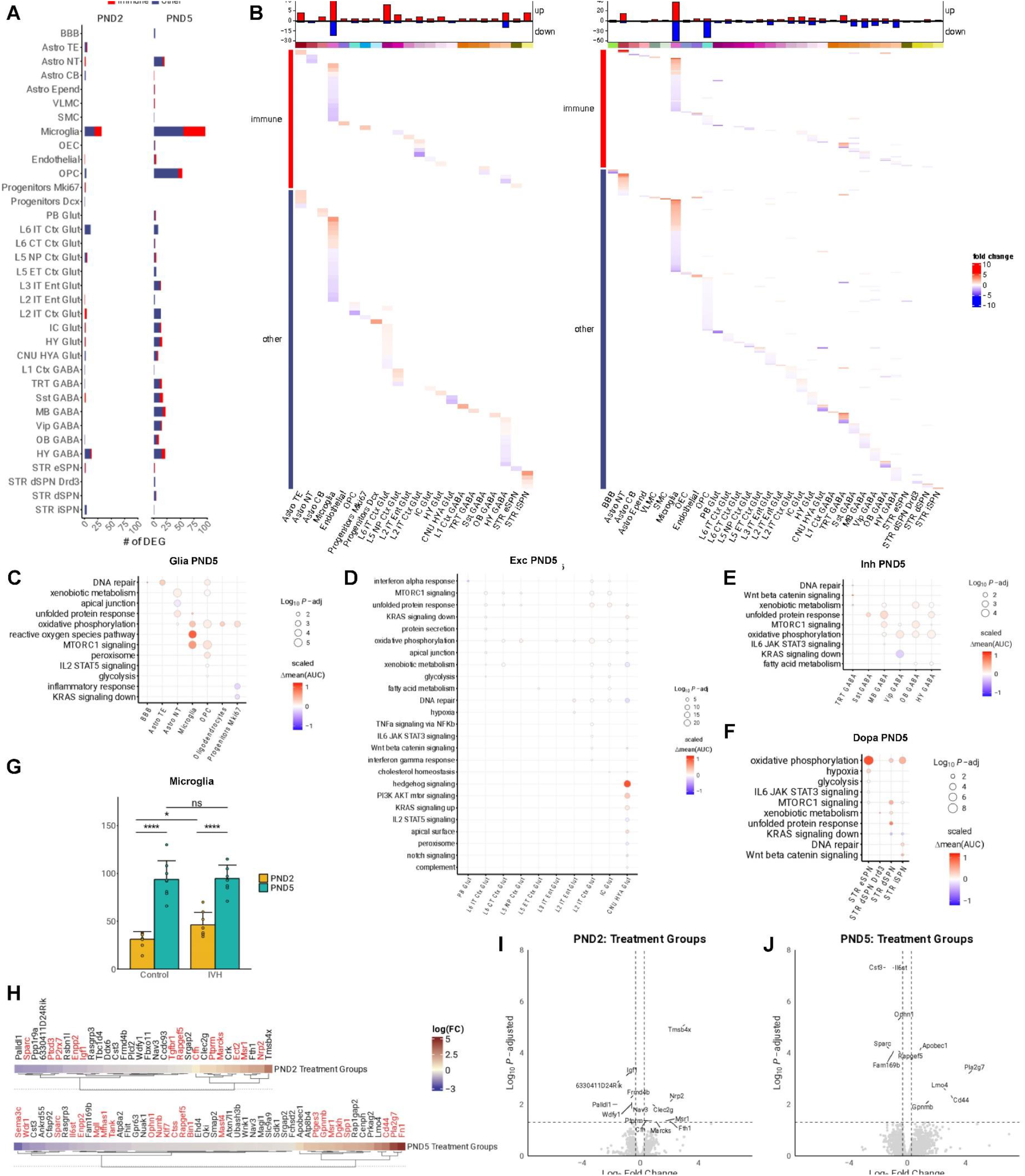
Control vs. IVH by developmental stage in glial cells. **A**. Stacked barplot showing the number of differentially expressed genes per cell type. Genes highlighted in red are involved in immune pathways. **B**. Heatmap visualizing the differentially expressed genes in each cell type ranked by fold change differences between IVH-treated and control rats. Fold changes indicate whether gene expression is higher or lower in IVH compared with control. **C-F**. Significant immune pathways enriched after IVH for each cell type. Color shows AUC values ranked by higher (red) or lower (blue) in IVH compared with control. There was no significant immune pathway enrichment for PND2. **G**. Barplot with standard errors depicting microglia proportions in IVH and Control at both time points. Stars correspond to the significance of changes (Wilcoxon Rank Sum test; **** = p < 0.001, * = p < 0.05). **H**. Heatmaps showing fold changes between control and IVH-induced rats for PND2 and PND5 in microglia. The genes shaded blue show statistically significant reduced effect size after IVH inducement as compared to control, while those shaded red show statistically significant increased effect size. Genes labeled in red are involved in immune pathways. **I-J**. Volcano plots highlighting the transcriptomic differences in microglial at PND2 (I) and PND5 (J). Y-axis corresponds to the -log_10_(FDR) and X-axis corresponds to the log_2_(Fold Change).

### Developmental Stage-Specific Signaling Reveals a Shift from Neuroinflammatory pathways to Complex Regulatory Phenotype in Microglia Following IVH

In IVH, key signaling pathways involving proteins upregulated in glial-to-microglial communication are predominantly associated with microglial phagocytosis and polarization toward an M1-like proinflammatory state. In contrast, neuronal signaling to microglia appears to engage pathways suppressing inflammatory activity and promote a reparative M2 microglial phenotype. These pathways were variable based on developmental stage (Figure 4). The ligands and receptors prioritized; that is, with higher impact and signaling in IVH than in the control, at PND5 were more associated with the Wnt signaling pathway in cascades promoting phagocytosis and demyelination (Figure 4A-B), and the top IVH-associated regulons, while largely overlapping, also showed differences between the timepoints (Figure 4C-D).

**Figure 4.**
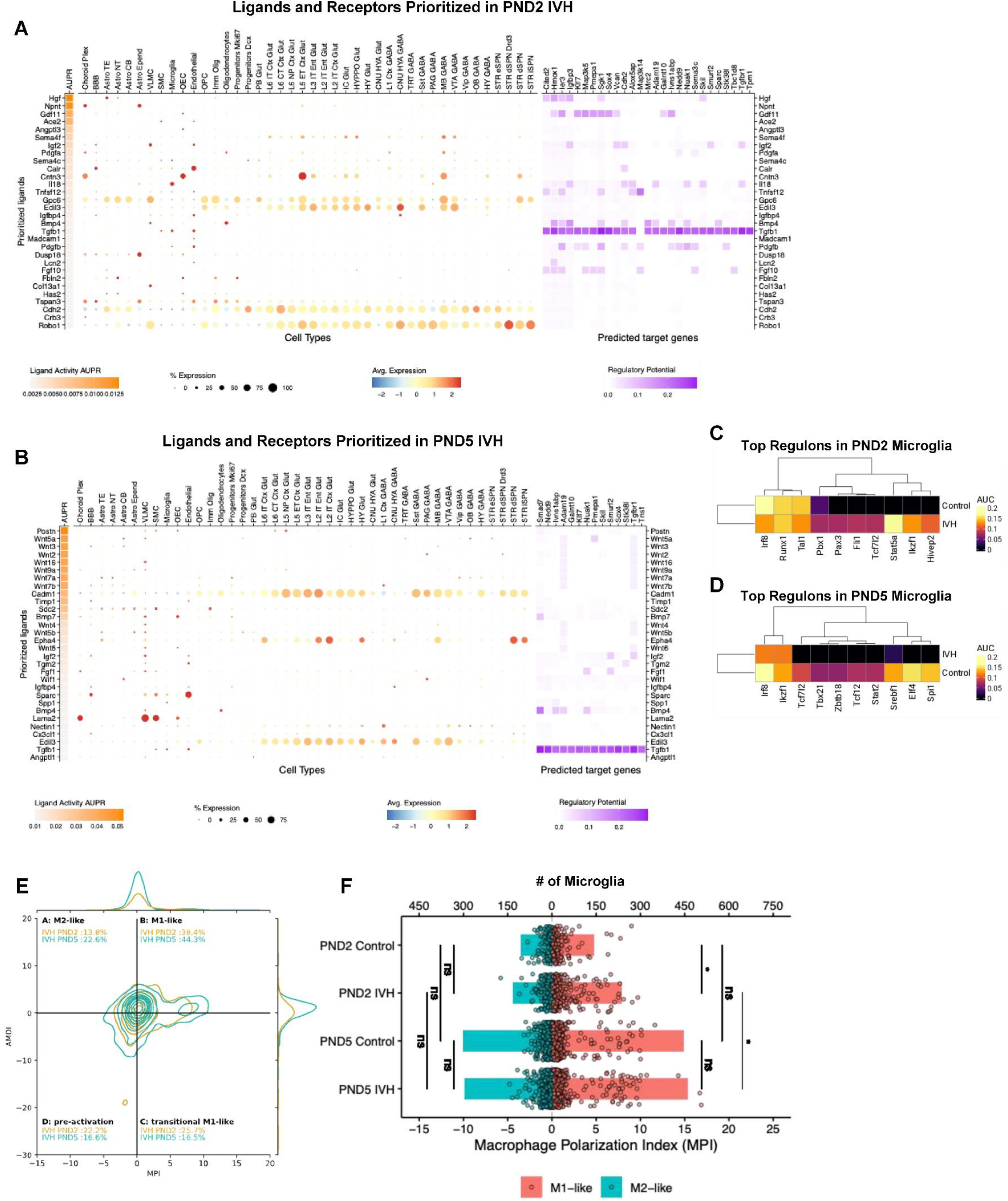
Control vs. IVH pathway enrichment and signaling changes in Microglia. **A-B**. Bubblechart showing the prioritized ligands after IVH for each cell type interactions with microglia. The heatmap on the left depicts the AUPR level, the size of the circle represents the percentage of cells expressing highlighted genes, and the heatmap on the right shows the predicted target genes and regulatory potential for each ligand at time point PND2 (A) and PND5 (B). **C-D**. Top 10 regulons ranked by AUC for microglia after IVH at time point PND2 (C) and PND5 (D). **E**. MacSpectrum plots of microglia states in IVH for PND2 and PND5 with relative predicted percentages. AMDI, Activation-induced Macrophage Differentiation Index; MPI, Macrophage Polarization Index. **F**. Mirrored barplot showing the macrophage polarization index (MPI) in IVH and Control for PND2 and PND5. Highlighted the significance based on Wilcoxon Rank Sum Test (* = p < 0.05).

At PND2, the response to IVH had no specific specialized pathway that stands out, but the top ligand seen at this time point, hepatocyte growth factor (HGF) has been implicated in neuroinflammation after traumatic brain injury [38]. Conversely, Igf2, another highly ranked ligand, inhibits microglial overactivation, suppresses proinflammatory responses, and prevents TNF-mediated cell death (Figure 4A) [39, 40]. Additionally, high levels of TGF-beta signaling in microglia, primarily expressed in microglia and endothelial cells after IVH, were also present and suppress pro-inflammatory and phagocytic activity of microglia (Figure 4A) [41, 42]. At PND5, however, a distinct proinflammatory signature emerges, dominated by non-canonical Wnt signaling (Figure 4B). This pathway has been implicated in neuroinflammation and neuronal demyelination, particularly through Wnt5a, Wnt7a, Wnt7b, and Wnt9a [43, 44]. Specifically, proinflammatory Wnt signaling primarily involves the top Wnt proteins seen as ligands here: Wnt3, which can activate proinflammatory signaling cascades in microglia and Wnt5a, which has been seen to promote neuronal maturation along with inflammatory microglial behavior [45-47]. Also seen highly ranked at PND5, Wnt7a is the Wnt protein most associated with demyelination, working in conjunction with myelin debris to increase oligodendrogenesis along with its related protein Wnt7b, which has been found to act as a signal triggering cell death [48, 49]. In contrast to this, cell adhesion molecules Cadm1, which plays a key roles in the myelination of neurons and activation of protective microglial behaviors was seen to be highly expressed by neurons as in signaling with microglia, likely as a protective response to Wnt signaling (Figure 4B) [50, 51]. Igf2 and TGF-beta were also seen as key effective ligands at PND5 but lower ranked and with less expression than at PND2 (Figure 4B).

### Transcriptional Regulators Underlying Microglial Polarization and Maturation After IVH

To identify key transcriptional drivers altered by IVH, we analyzed regulon activity in microglia and identified the top 10 IVH-associated regulons at each developmental timepoint (Figure 4C-D, Table S3). Runt-related transcription factor-1 (*Runx1*) was highly active in microglia at both PND2 and PND5, as described above. *Runx1* is a driver of microglial activation after CNS injury [52]. It is also known to modulate microglial phenotype via the Notch1 signaling pathway and interacts with immune-related pathways such as NF-κB and TLR4 [53, 54]. While *Runx1* activity showed minimal differences between IVH and control at PND2, its regulatory influence was markedly enhanced in IVH microglia at PND5, suggesting a time-specific activity during microglial reprogramming.

We also observed significant upregulation of *Stat5* regulon activity in IVH, a transcription factor central to cytokine signaling downstream of the JAK2–STAT5 pathway. *Stat5* has been proposed as a potential therapeutic target for neuroinflammation and is known to regulate CD44, a receptor we found to be significantly elevated in PND5 microglia following IVH (Figure 3H and J, Figure 4C) [55].

As many of the identified genes and regulons directly influence M1-and M2-like polarization states in microglia, we next assessed the extent and distribution of microglial polarization across developmental time points (Figure 4E-F). At PND5, the majority of microglia exhibited an M1-like proinflammatory phenotype (Figure 4E), but a larger proportion of cells had transitioned toward an M2-like reparative state compared to PND2, indicating maturation of polarization capacity. The macrophage polarization index (MPI), which measures inflammatory potential, of inflammatory microglia is significantly higher after IVH at PND5 compared to PND2 and in IVH at PND2 compared to PND2 control (Figure 4H). There was a trend toward increased polarization toward non-inflammatory phenotypes following IVH at both time points that did not reach statistical significance (Figure 4F).

Collectively, these data highlight that transcriptional regulation of microglial identity and function becomes more pronounced and specialized at PND5, with *Runx1* and *Stat5* emerging as key IVH-responsive regulators. This more mature transcriptional landscape is accompanied by both heightened proinflammatory signaling and increased capacity for reparative polarization, reflecting a dual-edged microglial response that may shape long-term outcomes following neonatal brain injury.

## DISCUSSION

This is the first report of how the neonatal brain’s transcriptomic reaction to injury differs between two time points in the neonatal period. Like our prior work, we have shown that the innate neuroimmune response is more profound after injury at PND5. However, these data reveal that there are changes in neuroimmune activation at PND2 that were not revealed by our prior flow cytometry analysis. This underscores the importance of utilizing high-resolution techniques such as RNAseq that better reveal cellular changes after injury.

IVH activated genes linked microglial phagocytic activity which can lead to neuronal injury, particularly in synaptic loss and demyelination. Interestingly, we identified anti-inflammatory signaling from neurons directed toward microglia, possibly as a neuroprotective mechanism after IVH. Specifically in microglia, our findings strongly suggest that HGF is likely a key driver of early inflammatory responses at PND2, with *Igf2* and TGF-β signaling functioning as counter-regulatory mechanisms (Figure 4). Furthermore, the developmental stage dramatically affected the microglial transcriptome, with the inflammatory response being far more specific to activation of inflammation and demyelination of neurons via Wnt signaling in the PND5 animals.

The most powerfully upregulated genes in glial cells were related to species and oxidative metabolism and reactive oxygen species. Our prior work exposing glial cells to blood breakdown products has shown powerful effects of blood products on cellular metabolism in oligodendrocytes and microglia [56, 57]. This widespread metabolic dysfunction points to metabolic function as a central switch that regulates the behavior of all cell types to IVH. The marked increase in reactive oxygen species and oxidative phosphorylation (OXPHOS) pathway genes in microglia at PND5 (Figure 3) suggests that microglia are key mediators of injury progression in response to IVH, in alignment with our previous findings [57]. This temporal shift indicates that as microglia mature during early postnatal development, their immune responses become increasingly robust, potentially exerting stage-specific effects on other neuronal cells.

Analysis of neuronal transcriptional profiles revealed strikingly heterogeneous responses among neuronal subtypes. Notably, Hedgehog signaling was upregulated in glutamatergic excitatory neurons, implying enhanced proliferation or differentiation in this cell type. This observation is consistent with a previous study showing that the Hedgehog pathway is activated in ischemic/hypoxic injury and promotes progenitor cell proliferation [58]. Conversely, dopaminergic neurons exhibited elevated expression of OXPHOS-related genes, which may reflect either a compensatory adaptation to cellular stress or a metabolic vulnerability exacerbated by IVH. These findings highlight a cell-type-specific reprogramming of neuronal metabolism, likely orchestrated by inflammatory cues, with activated microglia playing a central regulatory role.

There are several limitations of this study. First is the assessment of only one time point after IVH. Our prior work has shown that the spatiotemporal activation of immune cells evolves after IVH, therefore comparing only one time point does not complete the entire picture of the immune, neuronal or glial response to this lesion [33]. It is possible that as the innate neuroimmune system populates, the animals injected at PND2 develop a profile similar to those at PND5. The other limitation is that our data provide no spatial resolution of transcriptomic changes. Our prior work has shown that neuroimmune activation is concentrated in white matter after IVH, but that microglial presence in the developing gray matter is altered as a factor of time [33]. In this study, time had an effect on microglial numbers, but IVH did not. Therefore, it is possible that time-dependent effects in the gray matter overshadowed IVH-dependent effects in white matter. An additional limitation is that many computational tools and reference databases used for transcriptomic analysis are primarily designed and optimized for human or mouse datasets; this may reduce sensitivity to species-specific features in our model system, potentially interfering with relevant rat-specific biological signals. Future work will help us better understand this exact spatiotemporal profile of RNA changes after IVH.

In summary, these findings reveal a dynamic and developmentally regulated neuroimmune landscape in response to IVH, where the same injury triggers distinct cellular and molecular responses depending on the timing. By capturing these differences at single-cell resolution, our study highlights how maturation shapes microglial behavior, signaling networks, and metabolic responses, offering critical insights into how early-life brain injury may lead to divergent outcomes.

## Supporting information

Supplementary Table 1 Presto Markers

Supplementary Table 2 DEGs

Supplementary Table 3 AUC Scenic

## ACKNOWLEDGMENTS

We thank the technical support from the Cancer Prevention and Research Institute of Texas (CPRIT RP240610). We thank members of the Miller lab and Berto lab for their critical reading of the manuscript and helpful discussions.

## AUTHOR CONTRIBUTION STATEMENT

E.W-A. and M.Z. are the first authors; E.W-A. designed the analysis pipeline and performed the majority of the data analysis. M.Z. further optimized the analytical approach and wrote the initial draft of the manuscript. Both E.W-A. and M.Z. created several figures and participated in editing the manuscript. H.J.V., B.B.O. and K.P.U. contributed to data analysis, manuscript preparation and editing. S.D.O. contributed to experimental design, planning, and manuscript preparation and editing. B.A.M. conceived and supervised the project. S.B. supervised computational analysis and data interpretation.

B.A.M. and S.B. co-wrote the main manuscript text with E.W-A. and M.Z., incorporating feedback from all authors. All authors reviewed and approved the final manuscript.

## ETHICS, CONSENT TO PARTICIPATE, AND CONSENT TO PUBLISH

Declarations: not applicable.

## NOMENCLATURE

AMDI: Activation-induced Macrophage Differentiation Index
AUC: Area under the curve
BBB: Blood-brain barrier
CNS: Central nervous system
CPRIT: Cancer Prevention and Research Institute of Texas
CSF: Cerebrospinal fluid
CNU HYA: Glut Cerebral anterior hypothalamus glutamatergic neurons
DEGs: Differentially expressed genes
Dopa: Dopaminergic neurons
Exc: Excitatory neurons
FDR: False discovery rate
GEMs: Gel beads in emulsion
HGF: Hepatocyte growth factor
HEB: Hibernate E/B27/GlutaMAX medium
Inh: Inhibitory neurons
IVH: Intraventricular hemorrhage
k-NN: k-Nearest neighbors
MPI: Macrophage Polarization Index
OECs: Olfactory ensheathing cells
OPCs: Oligodendrocyte precursor cells
OXPHOS: Oxidative phosphorylation
PCs: Principal components
PND2: Postnatal day 2
PND5: Postnatal day 5
PCA: Principal component analysis
Runx1: Runt-related transcription factor-1
snRNA-seq: Single-nucleus RNA sequencing
UMAP: Uniform Manifold Approximation and Projection
UMIs: Unique molecular identifiers

