## Supplementery Figures for "Developmental Stage-Dependent Transcriptomic Responses to Neonatal Intraventricular Hemorrhage"

- Supplementary Figures 1-4
- Supplementary Tables
  - Table S1: Gene markers identified for each cluster.
  - Table S2: Pseudobulk Statistics: genes differentially expressed between IVH and Control in PND2 and PND5.
  - Table S3: AUC statistics for eRegulons for both IVH and Control at each time point.

A

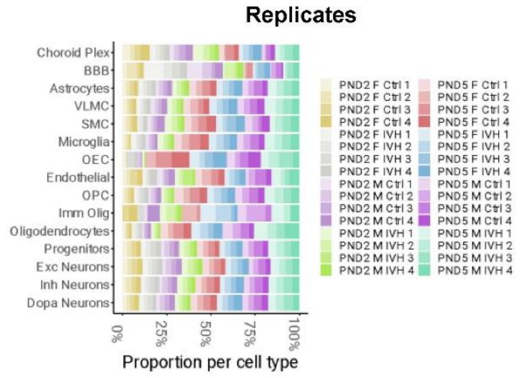


C

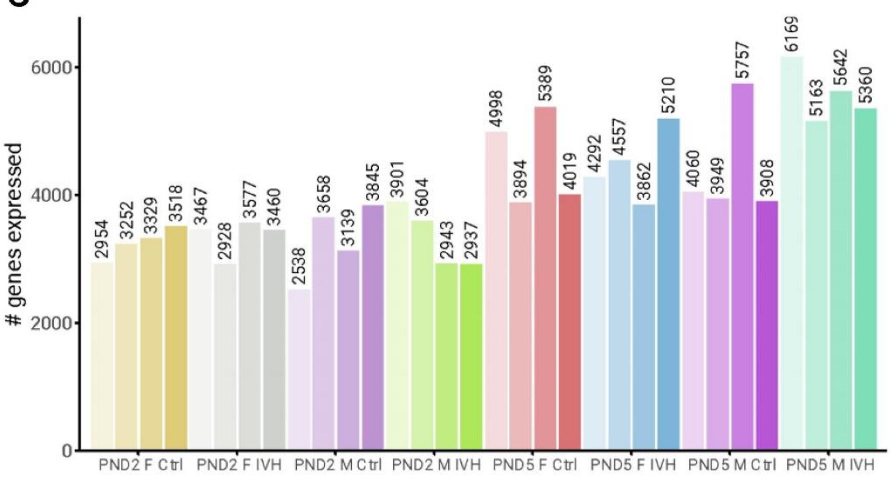


B

| | # nuclei | mean nGene | mean nUMI |
|---|---|---|---|
| *unfiltered* | 161933 | 1325 | 2740 |
| *filtered* | 129279 | 1409 | 2890 |

D

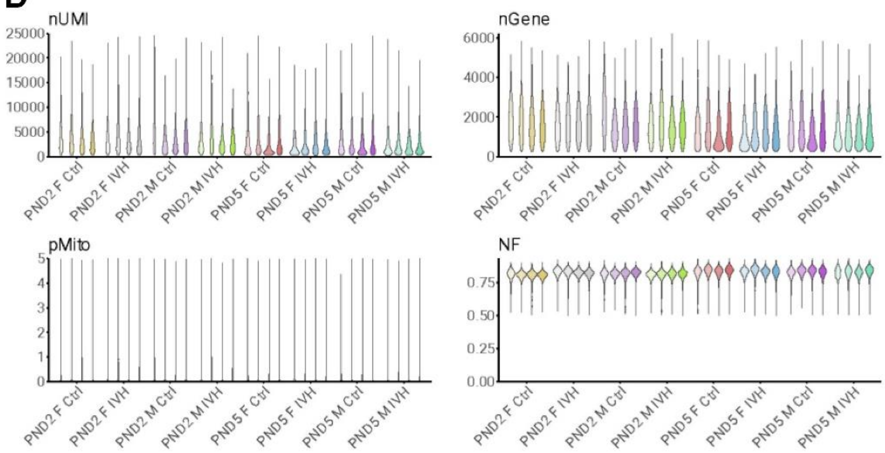


E

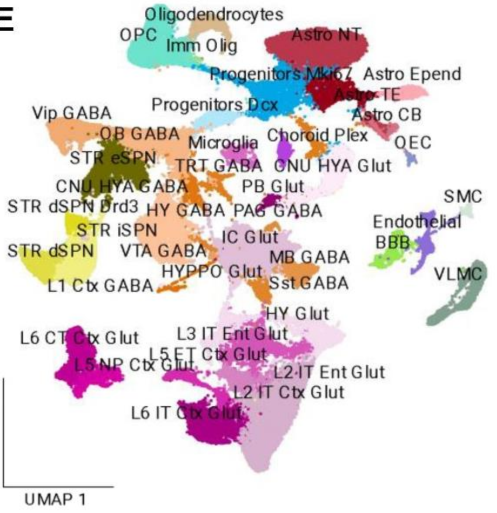


F

| | Markers: Excitatory Neurons |
|---|---|
| *PB Glut* | Lmx1a, Tll1 |
| *L6 IT Ctx Glut* | Il1rapl2, Hs3st2 |
| *L6 CT Ctx Glut* | Hs3st4, Slc17a7 |
| *L5 NP Ctx Glut* | Fezf2, Tshz2 |
| *L5 ET Ctx Glut* | Hs3st2, Slc17a7 |
| *L3 IT Ent Glut* | Ndst4, Scl17a7 |
| *L2 IT Ent Glut* | Rxfp1, Ntf3 |
| *L2 IT Ctx Glut* | Satb2, Glis3, Cux2, Hpse2 |
| *IC Glut* | Tfap2d, Maf |
| *HYPPO Glut* | Htr2c, Tmem91 |
| *HY Glut* | Sema3c, Scl17a6 |
| *CNU HYA Glut* | Slc17a6, Ebf1 |

G

| | Markers: Inhibitory Neurons |
|---|---|
| *L1 Ctx GABA* | Ndnf, Reln |
| *CNU HYA GABA* | Gad1, Ccbe1 |
| *TRT GABA* | Pax2, Tfap2b |
| *Sst GABA* | Sst, Ndst4 |
| *PAG GABA* | Nwd2, Trpm3 |
| *MB GABA* | Tcf7l2, Ntng1 |
| *VTA GABA* | Zfp804b, Chrm2 |
| *Vip GABA* | Vip, Adarb2 |
| *OB GABA* | Dlx1, Gad1 |
| *HY GABA* | Gad1, Eya4 |

H

| | Markers: Dopaminergic Neurons |
|---|---|
| *STR eSPN* | Grm8, Casz1 |
| *STR dSPN Drd3* | Drd3, Robo1 |
| *STR dSPN* | Drd1, Rarb |
| *STR iSPN* | Drd2, Penk |

I

| | Markers: Glial Cells |
|---|---|
| *Choroid Plex* | Ttr, Htr2c |
| *BBB* | S100b, Fth1 |
| *Astro TE* | Adgrv1, Veph1 |
| *Astro NT* | Gja1, Aqp4 |
| *Astro CB* | Pax3, Tnc |
| *Astro Epend* | Dnah12, Ccdc162 |
| *VLMC* | Ranbp3l, Cped1 |
| *SMC* | Ebf1, Prkg1 |
| *Microglia* | C1qa, Ptprc |
| *OEC* | Apod, Bnc2 |
| *Endothelial* | Flt1, Itm2a |
| *OPC* | Pdgfra, Cspg4 |
| *Imm Olig* | Bcas1, Myt1 |
| *Oligodendrocytes* | Mbp, Mobp |
| *Progenitors Mki67* | Mki67, Egfr |
| *Progenitors Dcx* | Dcx, Topo2a |

**Supplementary Figure 1. Quality Control Metrics and Filtering of Single-Nucleus RNA-seq Data. A.** Stacked barplot showing the proportion of total cells in each sample for each cell type identified. **B.** Table of the number of nuclei, mean number of genes before and after filtering, and mean number of unique molecular identifiers (UMI). Approximately 30,000 nuclei were filtered out because they did not pass quality control checks, and average genes and UMIs per nuclei were higher after filtering. **C.** Grouped barplot showing the number of genes expressed per sample (after filtering). Samples are grouped by replicates of conditions in the data. **D.** Quality control filtering of the nuclei for each sample. Nuclei were filtered based on number of unique molecular identifiers (less than 25000 to eliminate potential doublets), number of genes present (more than 250 to eliminate low quality nuclei), percentage of mitochondrial genes (less than 5% to ensure primarily nuclear data), and nuclear fraction (over 0.5 to ensure that the majority of the sample was actually from the cell nuclei). Cutoffs are marked with a dashed line. **E.** UMAP depicting the aggregated data with all cell annotation for all subtypes. **F-I.** Tables showing markers for cell types and subtypes by category of cell type.

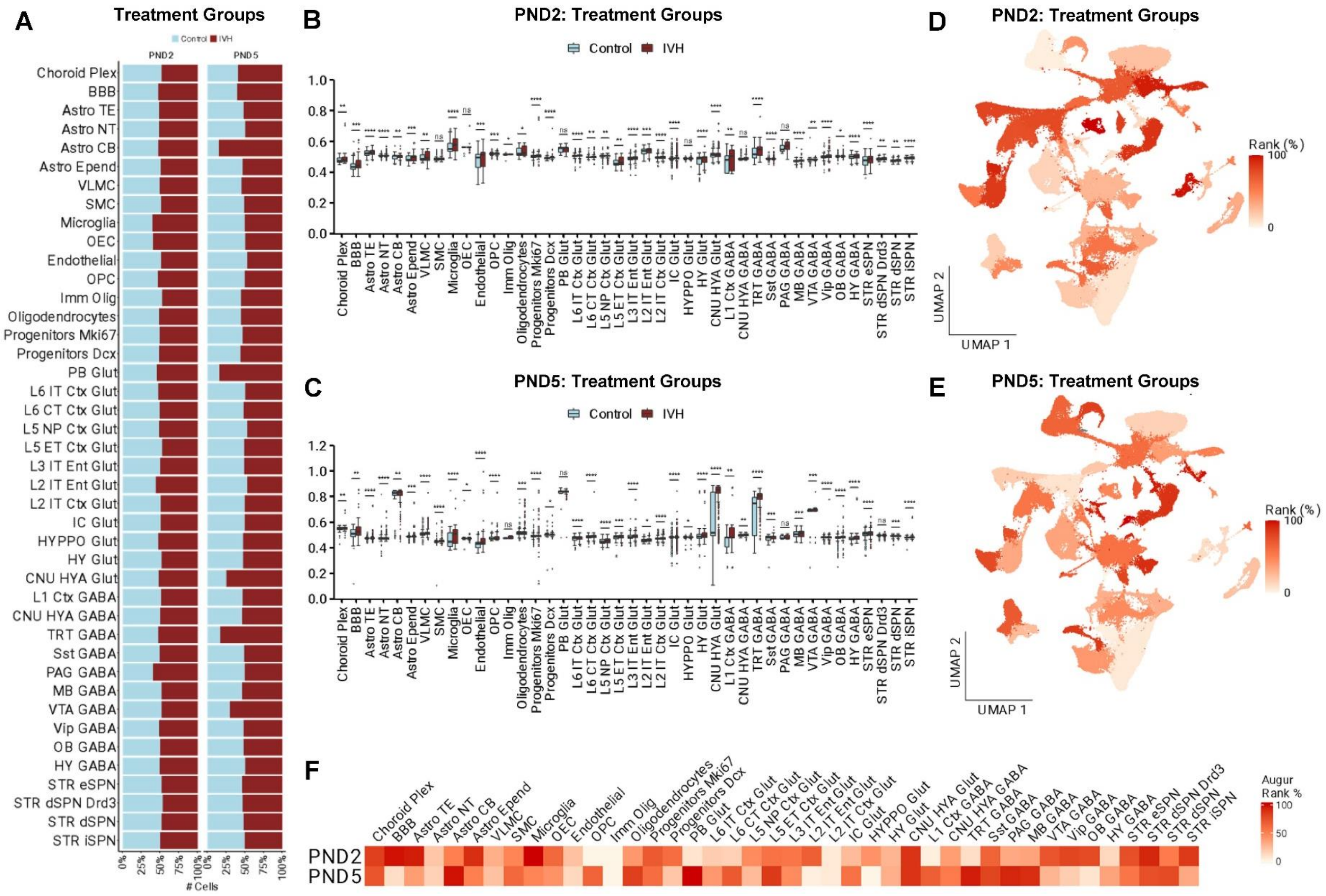


**Supplementary Figure 2. Cell-Type Specific Perturbation Signatures and Prioritization in Response to IVH at PND2 and PND5. A.** Stacked barplot showing the proportion of nuclei explained by IVH and control samples for each cell type, split by developmental time point. **B-C.** Grouped box and whisker plots showing MELD perturbation likelihood by cell type with statistical comparison between control and IVH at PND2 (B) and PND5 (C). Stars correspond to the significance of changes (Wilcoxon Rank Sum test; **** = $p < 0.001$, *** = $p < 0.005$, ** = $p < 0.01$, * = $p < 0.05$). **D-E.** UMAP of Augur cell prioritization ranked by effect of IVH treatment compared to control at PND2 (D) and PND5 (E). **F.** Heatmap of Augur ranks for each cell type based on effect of IVH treatment compared to control at PND2 and PND5.

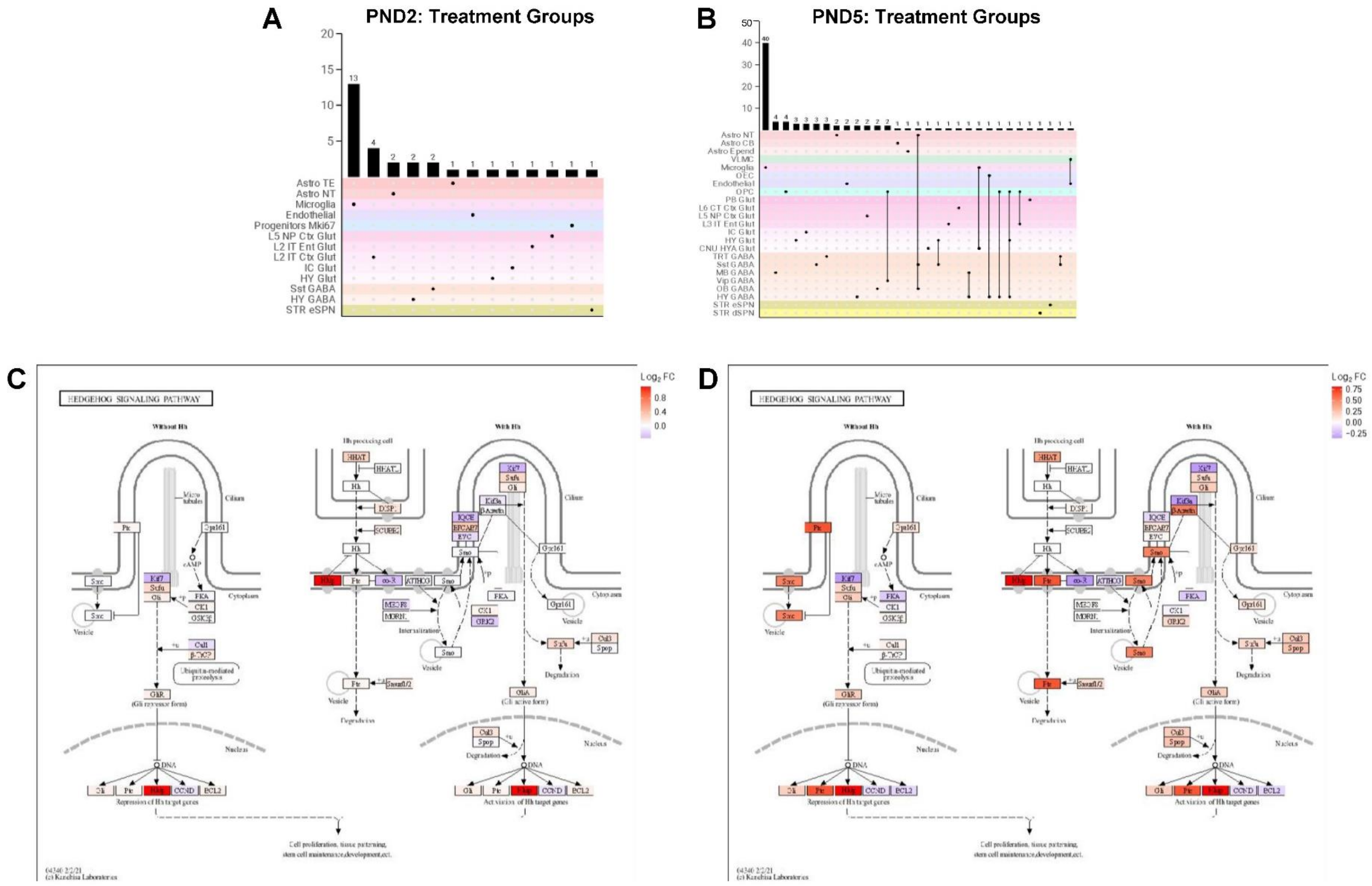


**Supplementary Figure 3. Developmental Stage–Specific Immune and Signaling Pathway Alterations in Response to IVH. A-B.** Upset plots showing numbers of overlapping differentially expressed genes involved in immune pathways at PND2 and PND5. Single dots with no connecting lines correspond to the bar in the bar plot marking the number of DEGs only significant in that cell type. Connected lines correspond to the bar in the bar plot marking the number of DEGs significant in both cell types. **C-D.** Hedgehog signaling KEGG pathway with pathway genes highlighted by log2(fold change) in cerebral anterior hypothalamus glutamatergic (CNU HYA Glut) neurons at PND2 (C) and PND5 (D). Genes are colored by fold change from control to IVH.

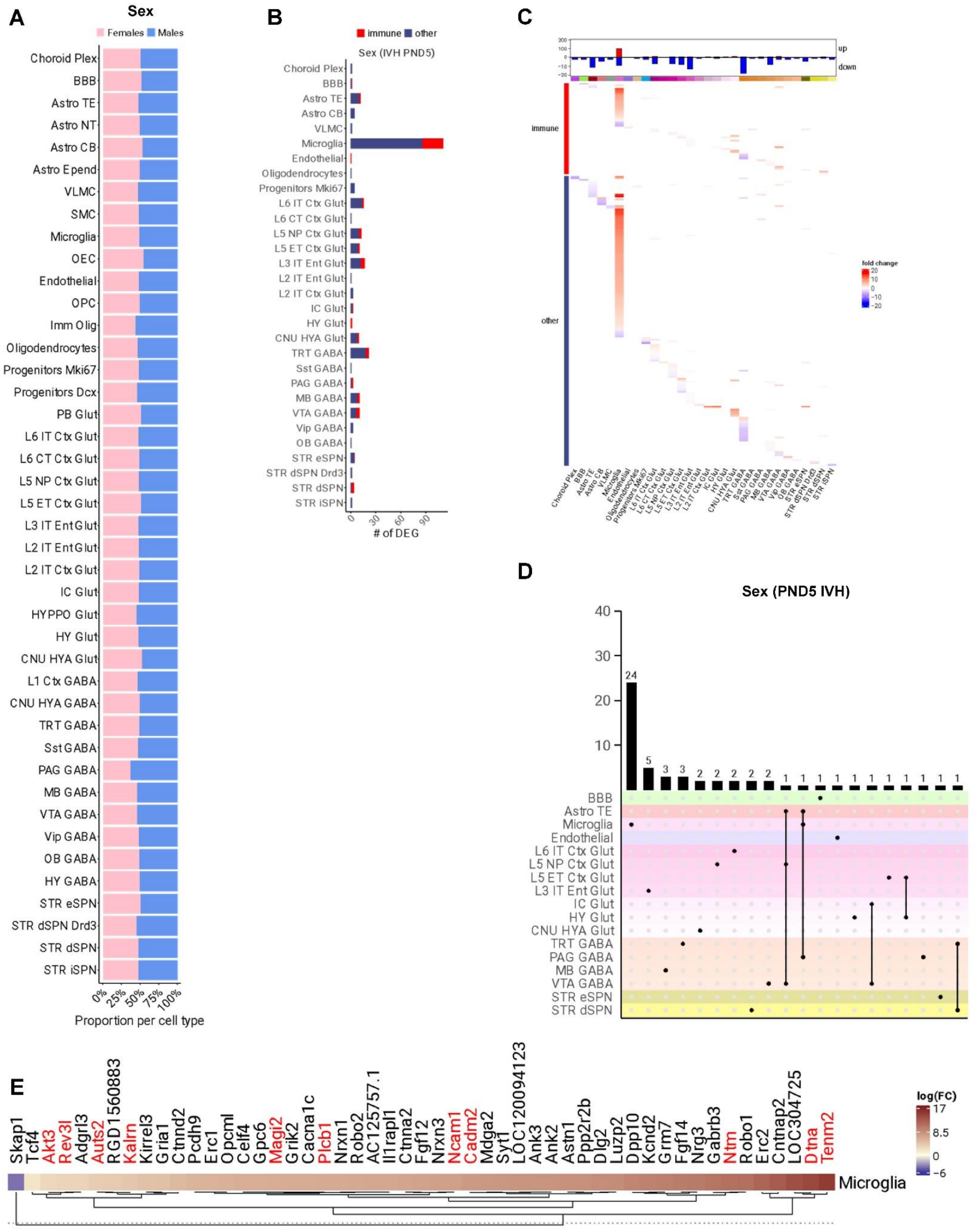

A
Sex
Females Males
Choroid Plex
BBB
Astro TE
Astro NT
Astro CB
Astro Epend
VLMC
SMC
Microglia
OEC
Endothelial
OPC
Imm Olig
Oligodendrocytes
Progenitors Mki67
Progenitors Dcx
PB Glut
L6 IT Ctx Glut
L6 CT Ctx Glut
L5 NP Ctx Glut
L5 ET Ctx Glut
L3 IT Ent Glut
L2 IT Ent Glut
L2 IT Ctx Glut
IC Glut
HYPPO Glut
HY Glut
CNU HYA Glut
L1 Ctx GABA
CNU HYA GABA
TRT GABA
Sst GABA
PAG GABA
MB GABA
VTA GABA
Vip GABA
OB GABA
HY GABA
STR eSPN
STR dSPN Drd3
STR dSPN
STR iSPN
0%
25%
50%
75%
100%
Proportion per cell type
B
immune other
Sex (IVH PND5)
### of DEG
0
30
60
90
C
up
down
immune
other
fold change
D
Sex (PND5 IVH)
40
30
20
10
24
5
3
3
2
2
2
2
2
1
1
1
1
1
1
1
1
1
1
E
Skap1
Tcf4
Akt3
Rev3l
Adgrl3
Auts2
RGD1560883
Kalrn
Kirrel3
Gria1
Ctnnd2
Pcdh9
Erc1
Opcml
Celf4
Gpc6
Magi2
Grik2
Cacna1c
Plcb1
Nrxn1
Robo2
AC125757.1
Il1rapl1
Ctnna2
Fgf12
Nrxn3
Ncam1
Cadm2
Mdga2
Syt1
LOC120094123
Ank3
Ank2
Astn1
Ppp2r2b
Dlg2
Luzp2
Dpp10
Kcnd2
Grm7
Fgf14
Nrg3
Gabrb3
Ntm
Robo1
Erc2
Cntnap2
LOC304725
Dtna
Tenm2
Microglia
log(FC)
17
8.5
0
−6

**Supplementary Figure 4. Glial cell sex-based changes in PND5 after IVH. A.** Stacked barplot showing the proportion of nuclei explained by male or female samples for each cell type. **B.** Stacked bar plot showing the number of differentially expressed genes per cell type. Genes highlighted in red are involved in immune pathways. **C.** Heatmap visualizing the differentially expressed genes in each cell type ranked by fold change differences between male and female rats. Fold changes indicate whether gene expression is higher or lower in males compared with females. **D.** Upset plot showing numbers of overlapping differentially expressed genes involved in immune pathways. Single dots with no connecting lines correspond to the bar in the bar plot marking the number of DEGs only significant in that cell type. Connected lines correspond to the bar in the bar plot marking the number of DEGs significant in both cell types. **E.** Heatmap showing fold changes between control and IVH-induced rats for PND2 and PND5 in microglia. The genes shaded blue show statistically significant reduced effect size in males as compared to females, while those shaded red show statistically significant increased effect size. Gene names in red are involved in immune pathways.
